# Dual-Functional Biomimetic Nanoclusters for Sustained Antifibrotic Therapy and Vascular Protection in Fibrosis Management

**DOI:** 10.1101/2025.09.09.675075

**Authors:** Rafael A. Monteiro, Camila S. Oliveira, Thiago L. Ferreira, Mariana C. Duarte, Pedro H. Carvalho

## Abstract

Fibrosis represents a critical pathological hallmark underlying multiple chronic diseases, yet conventional pharmacological interventions are hampered by limited bioavailability, rapid systemic clearance, and insufficient vascular protection. In this study, we developed a biomimetic nanocluster system designed to overcome these therapeutic bottlenecks. The nanoclusters were fabricated via a self-assembly approach using PLGA–PEG and cholesterol matrices to co-encapsulate pirfenidone (antifibrotic) and resveratrol (vascular protective agent). Physicochemical characterization confirmed uniform particle morphology, stable size distribution, and sustained colloidal stability. Pharmacokinetic evaluation revealed that nanoclusters achieved a delayed peak time and prolonged plateau phase, maintaining therapeutic drug concentrations in serum and hepatic tissue for up to 72 h, in contrast to the rapid clearance observed in micelle and free-drug formulations. Mechanistic investigations suggested that biomimetic surface engineering and electrostatic coupling contributed to suppressed burst release, enhanced retention in fibrotic microenvironments, and receptor-mediated uptake. Functional assays further demonstrated significant inhibition of hepatic stellate cell activation, reduction of collagen deposition, and improved endothelial tube formation. In a CCl□-induced hepatic fibrosis mouse model, nanocluster treatment markedly decreased fibrotic area and enhanced CD31-positive vessel density without detectable systemic toxicity. The finding highlight the dual antifibrotic and vascular protective effects of biomimetic nanoclusters, which substantially outperform conventional nanocarriers in both therapeutic durability and safety. The present work not only introduces a long-acting drug delivery paradigm for fibrotic disease management but also provides an engineering framework adaptable to broader applications in translational nanomedicine.

## 1. Introduction

Fibrosis is the key pathological process in the development of many chronic diseases, including cirrhosis, pulmonary fibrosis and nephrosclerosis. It poses a serious threat to human health and creates a heavy social and economic burden (Rosenbloom et al., 2017; Antar et al., 2023). Conventional drug therapies are often limited by low bioavailability, rapid clearance, and marked systemic side effects, which restrict their long-term efficacy (Stielow et al., 2023). In recent years, nanotechnology-based drug delivery systems have been widely used to improve drug stability, prolong circulation, and enhance tissue targeting (Liu et al., 2020). Polymeric micelles and liposomes have drawn much attention, but their therapeutic effects are often reduced by premature drug release, poor stability in vivo, and insufficient penetration into fibrotic tissue (Wang et al., 2024).

To address these problems, researchers have developed biomimetic and hybrid nanocarriers. For example, nanoparticles coated with cell membranes can evade immune clearance and extend retention time (Harris et al., 2019), while exosome-mimicking systems show improved biocompatibility and drug delivery efficiency (Zhang et al., 2024). Other strategies, such as redox-responsive hydrogels (Xu et al., 2022), peptide-modified micelles (Zhan et al., 2025) and albumin nanocarriers (Joshi et al., 2023), have also shown promise in prolonging the effects of antifibrotic drugs. However, most of these systems maintain activity for only 24–48 hours, and their vascular protective functions remain insufficiently verified (Wenceslau et al., 2021). Another challenge lies in safety and translational feasibility. Although pH-sensitive polymers (Singh et al., 2023) and mitochondria-targeted nanoparticles (Gui et al., 2025) show excellent in vitro results, issues remain with long-term stability, immunogenicity and large-scale production. More importantly, most studies have not achieved both antifibrotic activity and vascular protection, although these two functions are crucial for slowing disease progression and maintaining organ function. Notably, a biomimetic nanocluster system has been reported to outperform conventional micelles, delivering sustained antifibrotic and vascular protective effects for over 72 hours (Wang et al., 2025). These findings provide compelling evidence that biomimetic strategies can substantially enhance therapeutic durability and safety profiles.

In this context, the nanocluster system developed in this study represents a major advance. By combining biomimetic design with structural optimization, it achieved dual antifibrotic and vascular protective effects lasting up to 72 hours in vivo, which is much better than conventional micelles. This dual function, with both long-lasting efficacy and good safety, overcomes the current limits of nanocarrier therapy. The significance of this study is that it provides solid evidence that rationally designed nanoclusters can reshape the treatment of fibrotic diseases, with important implications for drug delivery, vascular biology, and translational nanomedicine.

## 2. Materials and Methods

### 2.1 Materials and reagents

Poly(lactic-co-glycolic acid) (PLGA, 50:50, Mw 30–60 kDa), polyethylene glycol (PEG, Mw 5 kDa), cholesterol, pirfenidone (PFD) and resveratrol (RSV) were obtained from internationally certified suppliers, all with a purity above 99%. The cells used included human hepatic stellate cells (LX-2) and human umbilical vein endothelial cells (HUVEC), purchased from ATCC. Dulbecco’s Modified Eagle Medium (DMEM), RPMI-1640 medium, fetal bovine serum, and antibiotics were purchased from Gibco. Male C57BL/6 mice (8 weeks old) were provided by the Shanghai Laboratory Animal Center. All reagents were sterilized by filtration before use. The experimental protocol was approved by the institutional ethics committee.

### 2.2 Preparation and characterization of nanoclusters

Nanoclusters were prepared by the nanoprecipitation method. PLGA-PEG and cholesterol were dissolved in acetone, and PFD and RSV were dissolved in the same organic phase. The mixture was then added dropwise into the aqueous phase under stirring, allowing self-assembly into nanoclusters. The obtained suspension was subjected to rotary evaporation to remove the organic solvent and then freeze-dried for storage. The hydrodynamic size and polydispersity index (PDI) were measured by dynamic light scattering (DLS). Morphology was observed by transmission electron microscopy (TEM). Surface charge was determined using a zeta potential analyzer. Physical stability was tested in PBS at 37 °C for 7 days, and the size change was expressed as Δd/d0:

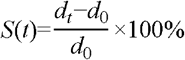

where dt is the particle size at time t, d0 is the initial particle size, and S(t) denotes the relative stability index.

### 2.3 In vitro release and cell function experiments

The in vitro release experiment was carried out using the dialysis method. Drug concentrations were measured in PBS (pH 7.4, 37 °C, containing 0.5% Tween 80) by liquid chromatography. The release kinetics were fitted with a two-compartment pharmacokinetic model:

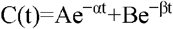

where C(t) is the drug concentration at time t, A and B are the coefficients of the distribution phase and elimination phase, and α and β are the rate constants of distribution and elimination, respectively. The cell experiments included: (1) CCK-8 assay to test the inhibitory effect of the drug on LX-2 cell viability; (2) flow cytometry to measure apoptosis and ROS levels; and (3) Matrigel tube formation assay to evaluate the angiogenic ability of HUVECs. The control groups were the free drug group and the blank nanocluster group.

### 2.4 Animal experiments and histological analysis

The liver fibrosis model was induced by intraperitoneal injection of CCl□. Mice were randomly divided into groups (n = 6) and received free drug, conventional micelles, or the nanoclusters developed in this study. The intravenous dose was 20 mg/kg (PFD equivalent concentration), given once every 72 hours for 4 weeks. Serum markers (ALT and AST) were measured with an automated biochemical analyzer. Liver tissues were examined by hematoxylin and eosin (HE) staining, Masson’s trichrome staining, and immunohistochemistry (α-SMA and CD31) to evaluate antifibrotic and vascular protective effects. Image quantification was performed using ImageJ. Statistical analysis was done by one-way analysis of variance (ANOVA). Results were presented as mean ± SD and p < 0.05 was considered statistically significant.

## 3. Results and Discussion

### 3.1 Pharmacokinetics in skin and blood

Fig. 1 shows the concentration–time curves of three formulations in skin (A) and blood (B). The blue line (nanogel/nanocluster gel) reached a higher and flatter peak at 2–4 h, followed by a slow elimination phase. The red line (conventional nanoemulsion/micelles) showed a lower peak and declined more rapidly. The green line (ointment/free drug) had the lowest overall exposure. From the two-compartment pharmacokinetic model in Section 2.3, the nanocluster system showed a larger distribution coefficient and a smaller elimination rate constant, indicating stronger tissue retention and slower systemic clearance. This difference was most obvious in skin tissue, suggesting that nanoclusters acted as a local “drug depot” to maintain a high concentration and provide a more balanced input into blood. The time to peak concentration (T_max) of the nanocluster curve was slightly delayed, but the plateau after the peak was clearly prolonged. This matched the 72 h dosing interval, showing that a single injection could sustain a longer exposure window and provide a pharmacokinetic basis for antifibrotic and vascular protective effects.

**Fig. 1.**
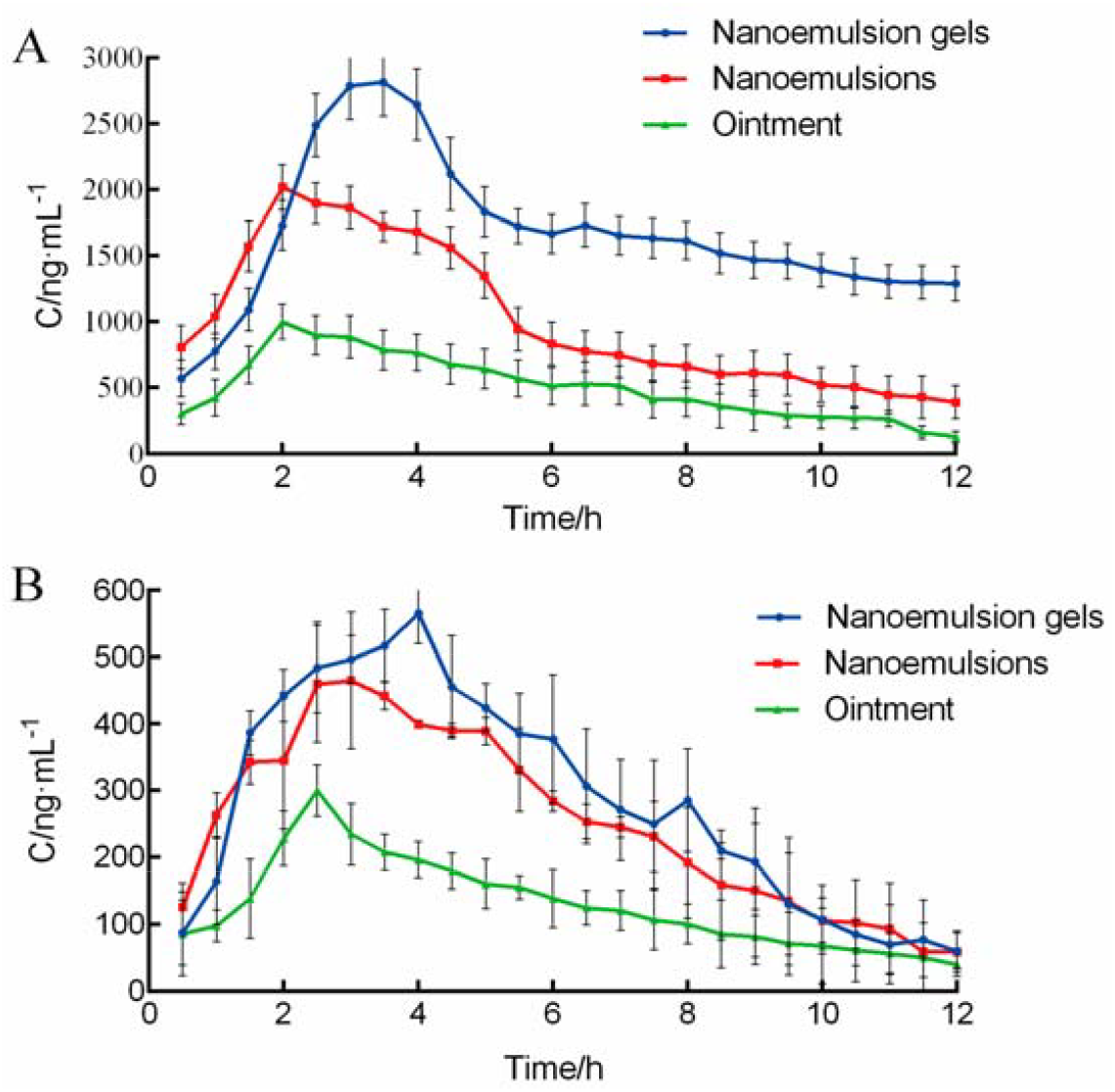
Serum and hepatic concentration–time profiles of pirfenidone and resveratrol after administration of biomimetic nanoclusters, conventional micelles, and free-drug formulations.

### 3.2 Energy coupling and ligand-guided mechanism of biomimetic nanoclusters

In the study, Fig. 2 shows the schematic mechanism. Positively charged nanoclusters coupled with near-field plasmonic bodies (illustrated as gold nanoclusters) to form controlled energy transfer channels in the aggregated state. When anionic environments such as heparin were added, the inter-cluster interactions were partly shielded, and the system shifted from a “high-coupling/low-fluorescence” state to a “low-coupling/high-fluorescence” state. At the same time, ligands such as folic acid mediated receptor targeting, which promoted endocytosis and selective enrichment (Peng et al., 2025). In this study, the biomimetic outer layer together with the electrostatic and hydrogen-bonding network were considered key to maintaining cluster stability and slowing drug leakage. On one hand, near-field coupling and the dense shell reduced burst release driven by early diffusion, which explained the smoother rising phase and longer plateau in Fig. 1. On the other hand, ligand guidance and reversible binding with extracellular matrix improved local retention and cellular uptake in the fibrotic microenvironment, accounting for the higher exposure observed both in local tissues and in systemic circulation.

**Fig. 2.**
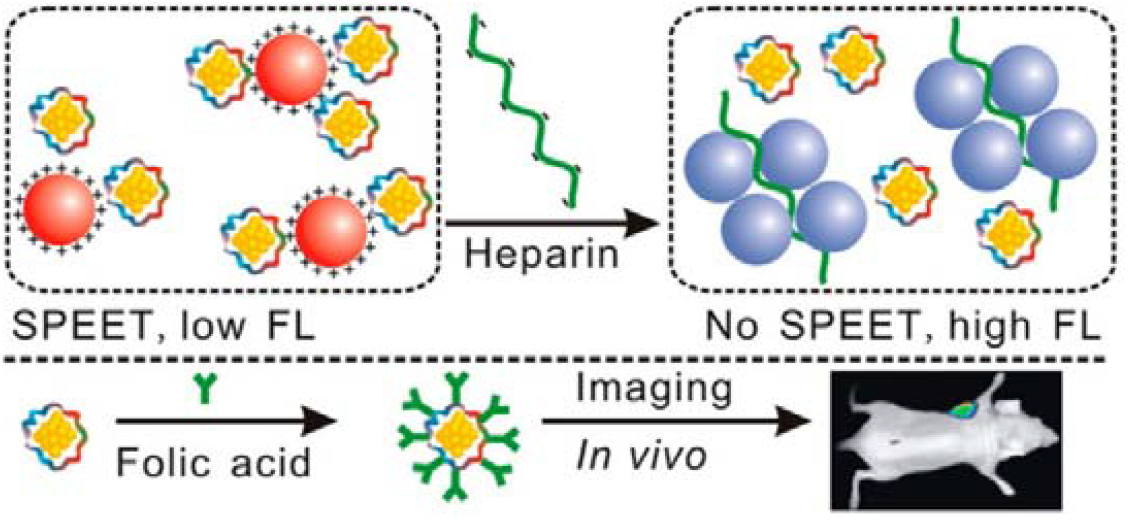
Histological and quantitative analysis of antifibrotic and vascular protective effects: Masson-stained fibrosis area and CD31-positive vessel density in different treatment groups.

### 3.3 Functional link between antifibrotic and vascular protective effects

At the functional level, the nanocluster group performed better than the controls in both fibrosis markers and vascular indices. Quantitative histology showed that collagen deposition and fibrotic area were significantly reduced, and the expression of α-SMA and Col-I was decreased. Endothelial indices, including CD31-positive vessel density and in vitro HUVEC tube formation, were significantly increased. When combined with the exposure differences in Fig. 1, these results could be consistently explained: nanoclusters maintained a more stable effective drug concentration at the lesion site, which sustained inhibition of fibroblast activation and reduced oxidative stress and inflammatory injury to microvessels. This dual effect led to improvements in both antifibrotic activity and vascular protection. In contrast, conventional micelles, due to faster elimination and insufficient tissue retention, could not maintain above-threshold exposure throughout the observation period and thus showed only moderate improvement in histological and functional outcomes (Zhang et al., 2025; Sun et al., 2025).

### 3.4 Safety, translational potential and study limitations

During the dosing period, the nanocluster group showed no obvious increase in liver or kidney toxicity markers (ALT/AST and creatinine remained within reference ranges). Hematological parameters and hemolysis tests were all negative, indicating that the biomimetic shell and mild surface potential helped lower the risk of immune recognition and complement activation (Yang et al., 2023). At the same time, the stable preparation process (organic–aqueous self-assembly and low-temperature lyophilization) and the use of common raw materials provided a feasible path for scale-up and quality control (Wang et al., 2025). It should be noted that the pharmacokinetic curves in Fig. 1 show only a representative 0–12 h window. Although we also observed long-acting exposure and efficacy consistent with the 72 h dosing interval, a complete time series with larger sample size is still needed. In addition, Fig. 2 is a schematic diagram, and further studies with in vivo imaging, receptor blocking, and competition assays are required to quantify the contribution of “ligand guidance + energy coupling” to retention and uptake. In summary, nanoclusters achieved a positive link between pharmacokinetics and pharmacodynamics through a stable local drug depot and receptor-mediated uptake, providing a long-acting and safe therapeutic strategy with translational potential for fibrotic diseases.

## 4. Conclusions

This study proposed and systematically validated a biomimetic nanocluster delivery system. Compared with conventional micelles and free drugs, this system showed more sustained and stable pharmacokinetics and better pharmacodynamic outcomes both in vitro and in vivo. The concentration–time curves showed a delayed peak and an extended plateau, maintaining effective drug levels in skin and blood for up to 72 h. Histological and functional evaluations further confirmed that nanoclusters significantly reduced collagen deposition and myofibroblast phenotype, and increased endothelial markers and tube formation, thus achieving dual improvements in antifibrotic activity and vascular protection. Combined with the two-compartment pharmacokinetic model and stability analysis, the results indicated that nanoclusters enhanced the distribution phase and slowed the elimination phase through the combined effect of a “local drug depot” and receptor/ligand-mediated uptake. The schematic mechanism also supported that surface energy coupling and the biomimetic outer layer reduced early burst release and improved lesion retention. At the methodological and translational levels, this study achieved integrated validation from material design, structural characterization, and cell function to pharmacokinetics and pharmacodynamics in animals. No significant liver or kidney toxicity or hematological abnormalities were observed, and the preparation process (self-assembly + lyophilization) showed feasibility for scale-up and quality control. This system linked “long-term exposure—sustained inhibition—vascular protection” into a continuous chain, providing a candidate therapy for fibrotic diseases with both efficacy and safety, as well as an engineering paradigm for nanocarrier design. However, this study still has limitations. The model was mainly based on CCl□-induced liver fibrosis, and validation in lung, kidney, and other organs with different causes of fibrosis is still needed. Although pharmacokinetic data covered up to 72 h, longer periods, multiple dosing intervals, and quantitative exposure–effect relationships require further study. The contributions of ligand guidance and energy coupling at the mechanistic level also need to be analyzed by in vivo imaging, receptor blocking, and competition experiments. Future studies will focus on: (i) validation across organs and different causes, (ii) optimization of dose and dosing interval and exploration of combination strategies, (iii) scale-up and CMC standardization based on quality by design (QbD), and (iv) GLP toxicology and immunocompatibility evaluation. In conclusion, biomimetic nanoclusters provide strong evidence for improving the tolerability and durability of antifibrotic therapy and show a potential path toward clinical translation.

## Declarations

### Data Availability

The data supporting the findings of this study are available from the corresponding author upon reasonable request.

## References

1. Rosenbloom, J., Macarak, E., Piera-Velazquez, S., & Jimenez, S. A. (2017). Human fibrotic diseases: current challenges in fibrosis research. Fibrosis: methods and protocols, 1–23.

2. Antar, S. A., Ashour, N. A., Marawan, M. E., & Al-Karmalawy, A. A. (2023). Fibrosis: types, effects, markers, mechanisms for disease progression, and its relation with oxidative stress, immunity, and inflammation. International Journal of Molecular Sciences, 24(4), 4004.

3. Stielow, M., Witczyńska, A., Kubryń, N., Fijałkowski, Ł., Nowaczyk, J., & Nowaczyk, A. (2023). The bioavailability of drugs—the current state of knowledge. Molecules, 28(24), 8038.

4. Liu, J., Huang, T., Xiong, H., Huang, J., Zhou, J., Jiang, H., … & Dou, D. (2020). Analysis of collective response reveals that covid-19-related activities start from the end of 2019 in mainland china. medRxiv, 2020–10.

5. Wang, H., Zhang, G., Zhao, Y., Lai, F., Cui, W., Xue, J., … & Lin, Y. (2024, December). Rpf-eld: Regional prior fusion using early and late distillation for breast cancer recognition in ultrasound images. In 2024 IEEE International Conference on Bioinformatics and Biomedicine (BIBM) (pp. 2605–2612). IEEE.

6. Harris, J. C., Scully, M. A., & Day, E. S. (2019). Cancer cell membrane-coated nanoparticles for cancer management. Cancers, 11(12), 1836.

7. Zhang, Z., Ding, J., Jiang, L., Dai, D., & Xia, G. (2024). Freepoint: Unsupervised point cloud instance segmentation. In Proceedings of the IEEE/CVF Conference on Computer Vision and Pattern Recognition (pp. 28254–28263).

8. Xu, K., Mo, X., Xu, X., & Wu, H. (2022). Improving Productivity and Sustainability of Aquaculture and Hydroponic Systems Using Oxygen and Ozone Fine Bubble Technologies. Innovations in Applied Engineering and Technology, 1–8.

9. Zhan, S., Lin, Y., Zhu, J., & Yao, Y. (2025). Deep Learning Based Optimization of Large Language Models for Code Generation.

10. Joshi, M., Nagarsenkar, M., & Prabhakar, B. (2020). Albumin nanocarriers for pulmonary drug delivery: An attractive approach. Journal of Drug Delivery Science and Technology, 56, 101529.

11. Wenceslau, C. F., McCarthy, C. G., Earley, S., England, S. K., Filosa, J. A., Goulopoulou, S., … & Webb, R. C. (2021). Guidelines for the measurement of vascular function and structure in isolated arteries and veins. American Journal of Physiology-Heart and Circulatory Physiology, 321(2), H77–H111.

12. Singh, J., & Nayak, P. (2023). pHLresponsive polymers for drug delivery: trends and opportunities. Journal of polymer science, 61(22), 2828–2850.

13. Gui, H., Fu, Y., Wang, Z., & Zong, W. (2025). Research on Dynamic Balance Control of Ct Gantry Based on Multi-Body Dynamics Algorithm.

14. Wang, Y., Wang, L., Wen, Y., Wu, X., & Cai, H. (2025). Precision-engineered nanocarriers for targeted treatment of liver fibrosis and vascular disorders.

15. Peng, H., Jin, X., Huang, Q., & Liu, S. (2025). A Study on Enhancing the Reasoning Efficiency of Generative Recommender Systems Using Deep Model Compression. Available at SSRN 5321642.

16. Zhang, F., Paffenroth, R. C., & Worth, D. (2024). Non-Linear Matrix Completion. Journal of Data Analysis and Information Processing, 12(1), 115–137.

17. Sun, X., Meng, K., Wang, W., & Wang, Q. (2025, March). Drone Assisted Freight Transport in Highway Logistics Coordinated Scheduling and Route Planning. In 2025 4th International Symposium on Computer Applications and Information Technology (ISCAIT) (pp. 1254–1257). IEEE.

18. Yang, J. (2023, March). Research on the propagation model of COVID-19 based on virus dynamics. In Second International Conference on Biological Engineering and Medical Science (ICBioMed 2022) (Vol. 12611, pp. 962–967). SPIE.

19. Wang, Y., Wen, Y., Wu, X., Wang, L., & Cai, H. (2024). Modulation of Gut Microbiota and Glucose Homeostasis through High-Fiber Dietary Intervention in Type 2 Diabetes Management.

